# Analysis of markerless limb tracking reveals chronic and progressive motor deficits after a single closed head injury in mice

**DOI:** 10.1101/2021.08.04.455083

**Authors:** Siobhán Lawless, Craig Kelley, Elena Nikulina, David Havlicek, Peter J. Bergold

## Abstract

**Background:** Acute injury following brain trauma may evolve into a chronic and progressive disorder. Chronic consequences of TBI have been understudied, in part, due to the lack of robust behavioral changes that are delayed in onset as well as chronic and progressive. Assessment of the chronic consequences of TBI also must distinguish behavioral changes that arises due to age vs those that develop and evolve over time due to injury.

**Methods:** C57BL/6 mice receive single closed head injury (CHI) and are analyzed at 7DPI, 14DPI or 180DPI on neurological severity score, open field, rotarod, beam walk, and simple-complex wheel.

**Results:** In the center of open field, injured mice have a turn bias at 180 days post-injury (DPI) not present at 7DPI. On rotarod, injured mice have shorter latencies at 7DPI, but not at 180DPI due to a large age effect in sham-injured mice. On beam walk at 180DPI, both sham and injured groups more slowly traverse a 2cm and 1cm beam than at 7DPI. Foot-faults show no significant effects of age or injury. On simple wheel injury affects speed at 14DPI with no effect on distance travelled. The lack of injury-dependent effects on beam walk or simple-complex wheel despite visible impairment was the impetus to assess limb position using Deeplabcut™ markerless tracking. Custom Python scripts were then developed to compute beam walk absition or foot fault severity (integral of limb displacement over time), and step frequency and quadrupedal limb coordination in simple-complex wheel. On the 2cm beam, age increased absition in all limbs of uninjured mice and both forelimbs of injured mice. On the 1cm beam both forelimbs and the left hindlimb of injured mice at 180DPI have larger absition than uninjured mice at 180DPI or injured mice at 7DPI. On a simple wheel injury affected speed only at 14 DPI with no effect on distance travelled. In contrast, injured mice at 180DPI developed a compensatory running strategy by increasing step frequency variability. This allowed injured mice at 180 DPI to reach sham-level quadrupedal limb coordination and improve running speed as compared to 14 DPI assessment. On complex wheel, injured mice at 180DPI do not express this compensatory running strategy resulting in impaired quadrupedal limb coordination. These data suggest chronic and progressive motor deficits of injured mice at 180DPI.

**Conclusions:** A single impact produces chronic and progressive motor deficits. Quantitative motor analysis using DeepLabCut™ tracking reveals deficits not seen using standard outcomes.

## Introduction

Moderate to severe clinical TBI produces chronic and progressive cognitive and motor deficits in one-third of patients with moderate to severe TBI(Foundas, 2013; Walker & Pickett, 2007; Wilson et al., 2017). Difficulties with balance, coordination, concentration, and information processing are common shortly after injury. Some individuals may fully recover while others have deficits that plateau and then worsen over time(Dean & Sterr, 2013; Johansson et al., 2009; Lundqvist et al., 2008; Niechwiej-Szwedo et al., 2007; Walker & Pickett, 2007). These two populations differ in the extent of radiological abnormalities as is evidenced by diffuse white matter injury(Kumar et al., 2009; Niogi et al., 2008; Wilson et al., 2017). Tasks requiring higher motor demand more easily detect deficits after clinical TBI(Vasudevan et al., 2014; Walker & Pickett, 2007). Patients with motor deficits may adopt compensatory suboptimal strategies to perform complex tasks(Martini et al., 2011; Vasudevan et al., 2014). In contrast, chronic and progressive motor deficits are less commonly seen in animal models of TBI as cognitive and motor deficits either appear to resolve or appear to remain unchanged over time(Shultz et al., 2020; Xiong et al., 2013). In a rat controlled cortical impact model, severe damage to the somatosensory cortex leads to long-lasting deficits in fine, but not gross, motor tasks(Ajao et al., 2012; Neumann et al., 2009). A challenge in studying chronic deficits in rodents is discriminating between motor decline that arises due to aging rather than to injury(Pischiutta et al., 2018). A second challenge is detecting compensatory behaviors that may conceal underlying deficits(Neumann et al., 2009; Vasudevan et al., 2014).

This study uses a mouse closed head (CHI) model of TBI in which the freely moving head of the mouse is impacted with an electromagnetically controlled piston that produces a contusion of the motor, somatosensory, association, and visual cortices as well as producing diffuse acute and chronic gray and white matter injury(Grin’kina et al., 2016; Havlicek et al., 2023). Aged injured mice utilized in this study develop progressive contralesional white matter atrophy and increasing density of phospho-tau expressing oligodendrocytes in subcortical regions such as thalamus and corpus callosum(Havlicek et al., 2023). CHI produces transient motor deficits, but it remains unknown if these deficits either persist or become chronic. We therefore examined whether deficits arise on Neurological Severity Score (NSS), open field, rotarod, beam walk, and simple-complex wheel in the subacute phase after injury, and whether or not they become progressive and chronic.

NSS qualitatively evaluates ten motor and behavioral tasks outcomes(Flierl et al., 2009). Open field assesses general locomotor activity, willingness to explore, basal anxiety and turning bias(Van Meer & Raber, 2005). The accelerating rotarod measures gait coordination, balance, and stamina(Schönfeld et al., 2017). While NSS qualitatively assesses complete beam traversal during beam walk, additional quantitative beam walk assessments include time to traverse and number of foot-faults(Carter et al., 2001; Luong et al., 2011; Schönfeld et al., 2017).

Simple/Complex wheel is a corpus callosum-dependent task that compares voluntary running on a simple wheel with uniformly spaced rungs vs running on a complex wheel with a complex rung pattern(Bishop et al., 1996; Hibbits et al., 2009; McKenzie et al., 2014). Assessment of running speed and distance travelled are used as surrogate markers of gait coordination. These standard task outcomes, however, do not take into account a role for suboptimal, compensating strategies in beam walk or simple/complex wheel(Flierl et al., 2009; Fox et al., 1998; Tsenter et al., 2008). Therefore, DeepLabCutTM was used as it employs deep learning to customize tracking the position of multiple sites on a mouse without physical tags(Mathis et al., 2018). Custom python scripts were developed to compute the fall time and displacement of each limb during a foot fault (absition) in beam walk; and step frequency, and limb coordination in simple-complex wheel. We report that assaying limb position detects compensatory, suboptimal strategies on beam walk and simple-complex wheel running that are not present in the subacute phase of injury.

## Methods

### Closed Head Injury (CHI) model of TBI

Male C57/BL6 mice (16 to 18 weeks, 26-28gr, Jackson Laboratories, Bar Harbor, ME) are randomly assigned to two groups and sham-CHI or CHI administered as previously published. Sham-CHI mice receive identical treatment as CHI mice without the impact. Mice are weighed on the day of injury and at 7 and 180DPI. Mouse weight increases between 7DPI and 180DPI (F_1,22_ = 26.7, p<0.0001). Mice at 180DPI (35.6±2.1g) are heavier than at 7DPI (30.9 ±2.2g) in both the sham-CHI and CHI groups (F_3,22_ = 10.8, p=0.0001, sham, p=0.0005, p=0.05). Mouse weight is neither affected by injury (F_1,22_ = 1.0, p=0.3) nor the interaction of injury and age (injury*age) (F_1,22_= 0.6, p=0.4).

After behavioral assessments, the brains of all mice undergo transcardial paraformaldehyde perfusion and the brains examined. Injured mice were included in behavioral assessments if the contusion site is centered 1.5mm lateral from the midline and -2mm posterior from Bregma. Injury site location excluded eight injured mice (7DPI, N=2; 180DPI, N=6). Two sham-CHI mice are excluded because their weight at 180DPI was greater than 2 standard deviations above the mean weight at that age (7DPI, 30.9 ± 2.2g; 180DPI, 35.6 ± 2.1g). All animal experiments are carried out in accordance with the NIH guide for the care and use of Laboratory animals and approved by the Institutional Animal Care and Use Committee of the SUNY-Downstate Medical Center (protocol #18-10558).

### Behavioral studies

Mice receiving sham-CHI or CHI received behavioral testing ending at 7, 14, or 180DPI (Sham-CHI 7DPI, N=10; CHI 7DPI, N=6; Sham-CHI 14DPI, N=4; CHI 14DPI, N=4; Sham-CHI 180DPI, N=6; CHI 180DPI, N=4). Behavioral studies are done in a windowless room with an average illumination of 268 lux and temperature of 24°C. In Neurological Severity Score (NSS), one point is assigned if a mouse is impaired in completing a task, and zero points if a mouse shows no impairment(Flierl et al., 2009). Two NSS tasks, open field, and beam walk, are assessed further using more quantitative metrics. In open field, mice are placed for 2 minutes in a 30cm diameter arena containing a centrally-located, computer-generated 10.25cm diameter circle. AnyMazeTM (Stoelting, Wood Dale, IL) software analyzes total distance travelled, time spent in the arena central circle and absolute turn angle. Absolute turn angle sums changes in center angle of the mouse’s torso as they move freely around the arena. Clockwise movements are assigned positive values in degrees, anti-clockwise movements are assigned negative values in degrees. Rotarod (Harvard Apparatus, Holliston, MA) testing consists of 5-minutes of training/habituation followed by 4 trials with the rod accelerating from 4– 40 rpm over 10 minutes with a 15-minute intertrial interval. Total latency is assessed.

Beam walk beams are 35cm long, 3cm, 2cm, or 1cm wide and elevated 10cm. Mice are recorded traversing the beam from each direction using an iPhone11Pro placed 18 inches lateral from the beam midpoint at 60 frames per second with 1080p and later converted to 720p resolution to optimize *post-hoc* limb tracking. Recordings begin when a mouse is placed on one beam end and ends after a mouse traverses the beam with all four feet planted on the opposite beam end. Mice traverse each beam twice, once in each direction with time to traverse defined as the average of the two trials. A foot-fault is recorded if the innermost fore or hindlimb digit cannot grip the top of the beam edge as described in Fox et al(Fox et al., 1998). Forelimb and hindlimb foot faults are assayed separately.

### Simple and Complex Running Wheel

Simple Running Wheel is done in a clear plexiglass activity wheel chamber (Lafayette Instruments Model 80820FS) containing a running wheel with regularly spaced rungs (Model 80820RW) for 30 minutes each day for five non-consecutive days Complex Running Wheel is done 30 minutes each day for five non-consecutive days in the same activity wheel chamber as simple wheel except with a running wheel containing an unevenly spaced rung pattern (Model 80821S) as referenced in McKenzie et al(McKenzie et al., 2014). Mice are supplied food and water *ad libitum* for each trial. After each trial, the activity wheel chamber and wheel are sprayed with 70% ethanol and wiped dry.

The running wheel chamber is placed on an elevated platform 2 inches high above a table surface. An iPhone 11 pro is placed in a level stationary position away from the chamber providing a working distance of 40.5cm between the camera lens and the wheel side chamber wall. This working distance allows for 2/3 of the wheel to be clearly visible from the camera perspective. High speed videos are recorded at 240 frames per second (fps) with 2x zoom at 720p resolution. The camera exposure is increased to +1.3x to enhance the lighting and color distinction between mouse paws and wheel rungs. Recording begins once the mouse steps onto the wheel and ends when the mouse steps off the wheel. Time interacting as well as time running on the wheel are recorded per wheel interaction. Speed is calculated using a stopwatch to record the time (seconds) and number of revolutions per interaction. Distance traveled is calculated as the number of revolutions completed per day.

### Motor analysis using DeepLabCut™

#### Beam Walk

Beam walk videos are further analyzed for *post-hoc* pose tracking and estimation using DeepLabCutTM v.2.2.0.2(Mathis et al., 2018). Briefly, 2 stationary and 6 anatomical points, including the top and bottom of the beam, both fore- and hindlimbs, nose, and base of tail, are marked manually in 146 frames from 4 different videos. Dataset creation and labeling are created with DeepLabCut™ installed under an Anaconda 3 environment. DeepLabCut™ Google Colaboratory is used for network creation, training, evaluation, and analysis of novel videos. Initial network is trained for 291,800 iterations reaching a train error of 1.25 pixels and test error of 4.36 pixels. Outlying labels are extracted, manually corrected, and merged into a new dataset and trained for 316,000 iterations with a train error of 1.23 pixels and test error of 2.34 pixels. One hundred forty-four new videos of beam walking are analyzed, and the horizontal (X) and vertical (Y) coordinates for each body part for each frame recorded. Custom Python scripts are written for extracted limb position data analysis. DeepLabCut™ labeled frames are utilized to extract the vertical position of the top of the beam. Pixel to centimeter conversion is measured using the distance in pixels and centimeters between the two DeepLabCut™ labeled beam points. Custom Python scripts extracted limb positions during beam walk. A 5-frame moving average window filtered the limb positions. Foot-faults are defined as instances with a limb falling below the surface of the beam. Total foot-fault absition for each limb is defined as the integral of vertical limb displacement (centimeters*seconds) below the top of the beam over time during all foot-faults. The python analysis code used to determine absition is publicly available at https://github.com/suny-downstate-medical-center/CHIanalysis.

### Simple-Complex wheel

DeepLabCut™ v.2.2.0.2 was used to further analyze simple and complex wheel videos for *post-hoc* pose tracking and estimation on days 3 of simple and complex wheel running. A total of 14 points (2 stationary and 12 anatomical) are marked manually in 250 frames from 10 videos (25 frames per video) differing in illumination (2 videos at +0.0x exposure, 8 videos at +1.3x exposure) and wheel type (4 simple wheel, 6 complex wheel), and training day. The two stationary points are marked from within the chamber and include the wheel center and wheel border. Six anatomical points including both fore- and hindlimbs, nose, and base of tail, are marked manually from within the chamber. Dataset creation and labeling are created as stated previously. Initial network is trained for 399,900 iterations reaching a train error of 1.77 pixels and test error of 15.27 pixels. Outlying labels are extracted, manually corrected, and merged into a new dataset four additional times. A fifth final dataset is trained for 646,400 iterations with a train error of 2.34 pixels and test error of 6.48 pixels. New videos of simple and complex wheel are analyzed, and the horizontal (X) and vertical (Y) coordinates for the wheel and each body part for each frame recorded.

Custom Python scripts extracted limb positions during running epochs defined as uninterrupted periods when the mouse is actively walking and or running on the wheel. Running epochs were manually identified by the experimenter with DeepLabCut™ labeled frames utilized to extract the positions of all four limbs. Artifacts caused by the running wheel temporarily obscuring limbs are corrected automatically by removing transients in extracted limb positions where limb position changed >15 pixels/frame and using third order spline interpolation. A 20-frame moving average window is used to filter the limb positions. The relative distance between the center of the animal and the end of each limb is computed for each frame. This is used to compute basic statistics for each running epoch, like step frequency. Limb coordination is defined as the cross correlation between relative distances to the center of the body for opposite limbs (e.g., right forelimb and left hindlimb) for all running epochs. Pixel to centimeter conversion is measured using the distance in pixels and centimeters between the two DeepLabCut™ labeled wheel points. The average conversion rate is 0.02 centimeters per pixel. The Python code for analyzing beam-walk and running wheel data is publicly available at https://github.com/suny-downstate-medical-center/CHIanalysis.

### Statistics

Mann-Whitney U test analyzes Neurological Severity Score. Kruskal-Wallis test analyzes foot-faults with Dunn’s post-hoc test. Repeated measures ANOVA analyzes treatment and time effects on rotarod. Two-way ANOVA analyzes weight at testing, open field, beam walk, and simple-complex wheel. Statistically significant differences on two-way ANOVA are further analyzed by one-way ANOVA with Tukey’s post hoc test for pairwise comparisons. Pairwise comparisons between the Sham-CHI at 7DPI or 14DPI and CHI 180DPI groups are not considered biologically relevant since both age and injury status differed. In all tests, statistical significance is set at 0.05.

## Results

NSS of sham or CHI mice lacked significant effects of injury (U=78.0, p=0.4) or age (U=66.5, p=0.9) at 7DPI or at 180DPI. On open field testing, sham and CHI mice traveled equal distances suggesting similar motivation to ambulate and explore (Table 1). Time in the center circle had no effect of injury (F_1,22_=0.6, p=0.5) but had a significant effect of age (F_1,22_=8.5, p=0.01) and injury*age (F_1,22_=5.6, p=0.03) (Figure 1, Panel A). Injured mice spend significantly more time in the center circle at 180DPI than 7DPI (F_3,22_=3.9, p=0.02, post-hoc, p=0.013). Absolute turn angle over the entire arena lacked a significant effect of injury (F_1,22_=3.3, p=0.1), age (F_1,22_=0.7, p=0.4), or injury*age (F_1,22_=2.2, p=0.2). Within the center circle, however, absolute turn angle had an effect of injury (F_1,22_=6.0, p=0.02), age (F_1,22_=8.0, p=0.01), and injury*age (F_1,22_=6.0, p=0.02) (Figure 1, Panel B). Post hoc test reveal a significant group effect (F_3,22_=5.2, p=0.007) with CHI mice turning more towards the ipsilesional hemisphere at 180 than 7DPI (p=0.013) or compared to sham mice at 180DPI (p=0.02). These data suggest that injured mice developed a progressive unilateral turning bias toward the ipsilesional hemisphere between 7 and 180 DPI.

**Table 1.** Open Field and Beam Walk. Distance traveled shows no significant difference due to injury (F_1,22_=4.0, p=0.06), age (F_1,22_=2.4, p=0.14), or interaction between injury and age (F_1,22_=0.001, p=0.9). Absolute turn angle assessed in the entire arena shows no significant effect of injury (F_1,22_=3.3, p=0.1), age (F_1,22_=0.7, p=0.4), or interaction between injury and age (F_1,22_=2.2, p=0.2). Forelimb foot-faults do not significantly differ on 3cm or 2cm, but was significant on the 1cm beam (3cm Left, H(3)=2.7, p=0.5, Right, H(3)=3.3, p=0.3; 2cm Left, H(3)=2.3, p=0.5, Right, H(3)=5.5, p=0.1; 1cm Left, H(3)=1.5, p=0.7, Right, H(3)=9.4, p=0.02). Post-hoc tests reveal no significant difference between groups. Hindlimb foot-faults did not significantly differ on the 3cm beam, but did differ significantly on the 2cm and 1cm beams (3cm Left, H(3)=1.6, p=0.7, Right, H(3)=6.5, p=0.09; 2cm Left, H(3)=8.9, p=0.03, Right, H(3)=1.2, p=0.7; 1cm Left, H(3)=17.1, p=0.001, Right, H(3)=8.3, p=0.04). Post hoc tests reveal only a significant difference between Sham-CHI7 and CHI180 mice. Thus, forelimb or hindlimb foot-faults have no significant effects of side, age, or treatment.

| Behavioral test | Parameter | Sham-CHI7 |  | CHI7 |  | Sham-CHI180 |  | CHI180 |  |
| --- | --- | --- | --- | --- | --- | --- | --- | --- | --- |
| Open Field | Distance traveled (meters) | 3.5 ± 0.5 |  | 4.6 ± 0.6 |  | 2.7 ± 0.3 |  | 3.8 ± 0.5 |  |
|  | Absolute Turn Angle in the arena (degrees) | 9047.2 ± 761.3 |  | 9381.7 ± 886.9 |  | 6722.2 ± 798.2 |  | 10037.3 ± 1713.8 |  |
| 3cm Beam Walk | Time to traverse beam (seconds) | 7.9 ± 0.8 |  | 16.3 ± 5.2 |  | 11.6 ± 1.4 |  | 12.5 ± 2.9 |  |
|  | Forelimb foot-faults | Left | Right | Left | Right | Left | Right | Left | Right |
|  |  | 0.1 ± 0.1 | 0.0 ± 0.0 | 0.0 ± 0.0 | 0.0 ± 0.0 | 0.0 ± 0.0 | 0.0 ± 0.0 | 0.0 ± 0.0 | 0.0 ± 0.0 |
|  | Hindlimb foot-faults | 0.5 ± 0.3 | 0.5 ± 0.3 | 0.3 ± 0.2 | 1.0 ± 0.7 | 0.3 ± 0.3 | 0.3 ± 0.3 | 1.0 ± 0.7 | 0.3 ± 0.3 |
| 2cm Beam walk | Time to traverse beam (seconds) | 7.8 ± 0.5 |  | 11.0 ± 1.4 |  | 12.6 ± 1.1* |  | 14.0 ± 1.6 |  |
|  | Forelimb foot-faults | Left | Right | Left | Right | Left | Right | Left | Right |
|  |  | 0.0 ± 0.0 | 0.0 ± 0.0 | 0.3 ± 0.3 | 0.0 ± 0.0 | 0.2 ± 0.2 | 0.0 ± 0.0 | 0.5 ± 0.5 | 0.3 ± 0.3 |
|  | Hindlimb foot-faults | 0.1 ± 0.1 | 0.6 ± 0.3 | 1.2 ± 0.4 | 1.0 ± 0.45 | 0.3 ± 0.2 | 0.5 ± 0.2 | 1.8 ± 0.9 | 1.0 ± 0.4 |
| 1cm Beam walk | Time to traverse beam (seconds) | 9.0 ± 0.7 |  | 12.1 ± 1.6 |  | 13.4 ± 2.4 |  | 25.1 ± 8.4* |  |
|  | Forelimb foot-faults | Left | Right | Left | Right | Left | Right | Left | Right |
|  |  | 0.3 ± 0.2 | 0.1 ± 0.1 | 0.8 ± 0.5 | 0.3 ± 0.3 | 0.3 ± 0.2 | 1.0 ± 0.3 | 1.0 ± 0.8 | 1.0 ± 0.5 |
|  | Hindlimb foot-faults | 0.6 ± 0.3 | 0.9 ± 0.3 | 2.3 ± 0.6 | 2.5 ± 0.5 | 3.2 ± 0.7 | 1.5 ± 0.7 | 6.0 ± 0.7 | 3.3 ± 1.1 |

**Figure 1.**
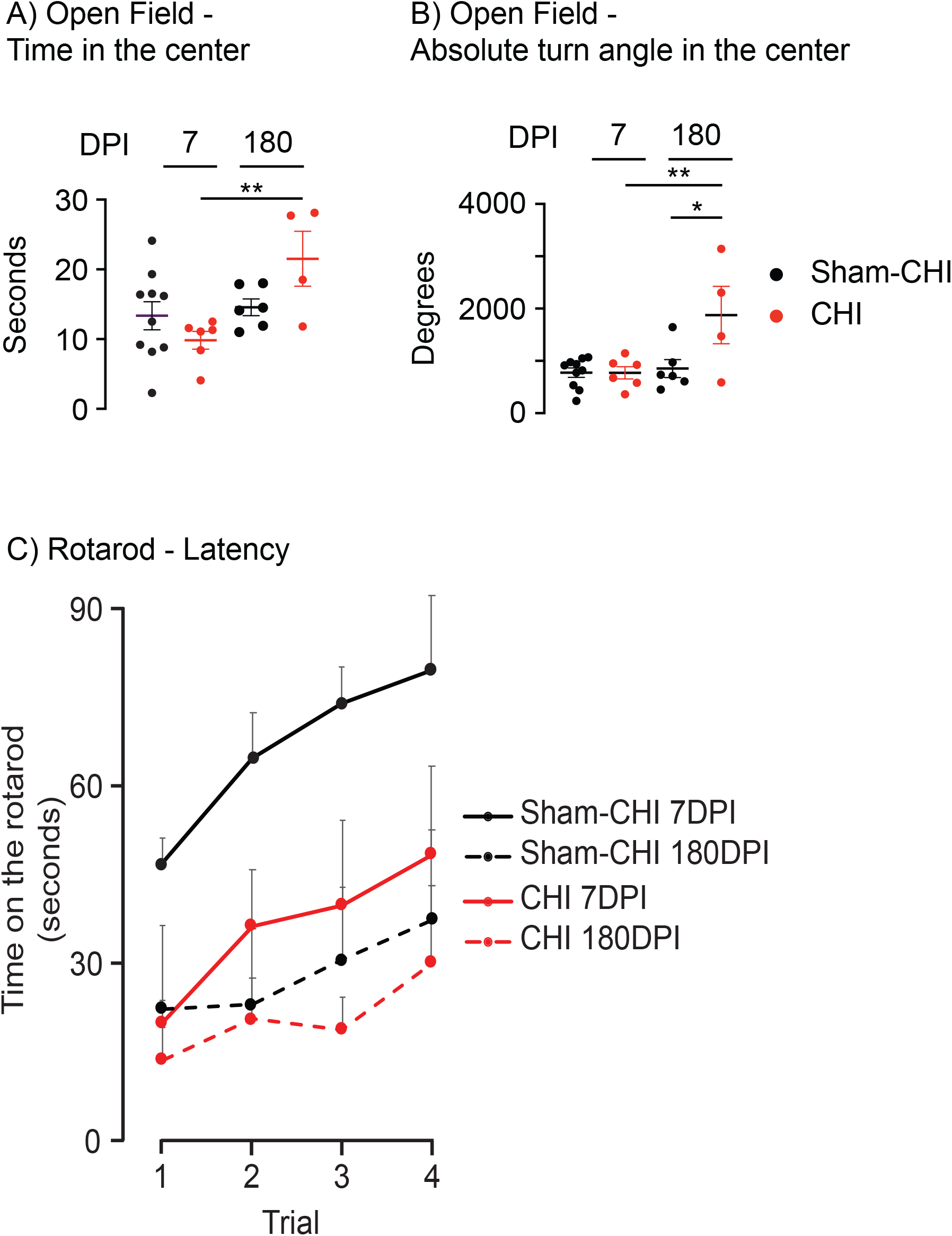
Open Field and Rotarod, **Panel A), Open Field - Time in the arena center** CHI mice differ significantly between 7 and 180 DPI (p=0.01**). **Panel B), Open Field - Absolute turn angle in the arena center**. CHI mice differ significantly between 7 and 180 DPI (p=0.01**). Sham-CHI and CHI mice differ significantly between at 180DPI (*p=0.02). **Panel C), Rotarod - Latency**, CHI and sham-CHI significantly differed on total rotarod latency at 7DPI but not at 180DPI (7DPI, F_1,14_=11.57, p=0.004**; 180DPI, F_1,7_=2.56, p=0.2). Latency of sham-CHI but not CHI mice differed between 7 and 180DPI (Sham-CHI, F_1,13_=5.96, p=0.03*; CHI, F_1,8_=0.49, p=0.51).

Injured mice at 7DPI have significantly lower latency to fall off the rotarod than sham mice (F_1,14_=11.6, p=0.004) (Figure 1, Panel C). In contrast, sham and CHI mice had similar latencies at 180 DPI (F_1,7_=2.6, p=0.2). Similar latencies at 180 DPI are due to a significant age effect that decreased latency between 7 and 180 DPI in sham mice (F_1,13_=6.0, p=0.03) but not injured mice (F1,8=0.5, p=0.5).

On beam walk, time to traverse the 3cm beam lacks significant effects of injury, age, and injury*age (Table 1, Table 2A). Time to traverse the 2cm beam has significant effects of injury, age but not injury*age. Time to traverse the 1cm beam also shows significant effects of injury, age but not injury*age. Biologically insignificant comparisons on the 2cm and 1cm beams are responsible for the injury effects. Sham-CHI mice have an age effect on the 2cm beam (p=0.01), and CHI mice have an age effect on the 1cm beam (p=0.05) (Table 2A). Foot-faults have no effect of injury or age on the 3cm, 2cm, or 1cm beams (Table 1). A simple foot-fault count, however, does not consider differences in the duration or distance the limb is off the beam that is assessed when measuring absition (Supplementary Video 1). Therefore, limb absition is measured on 3, 2, and 1cm beams providing additional information of foot fault severity not previously captured by counting the number of foot faults.

**Table 2.**
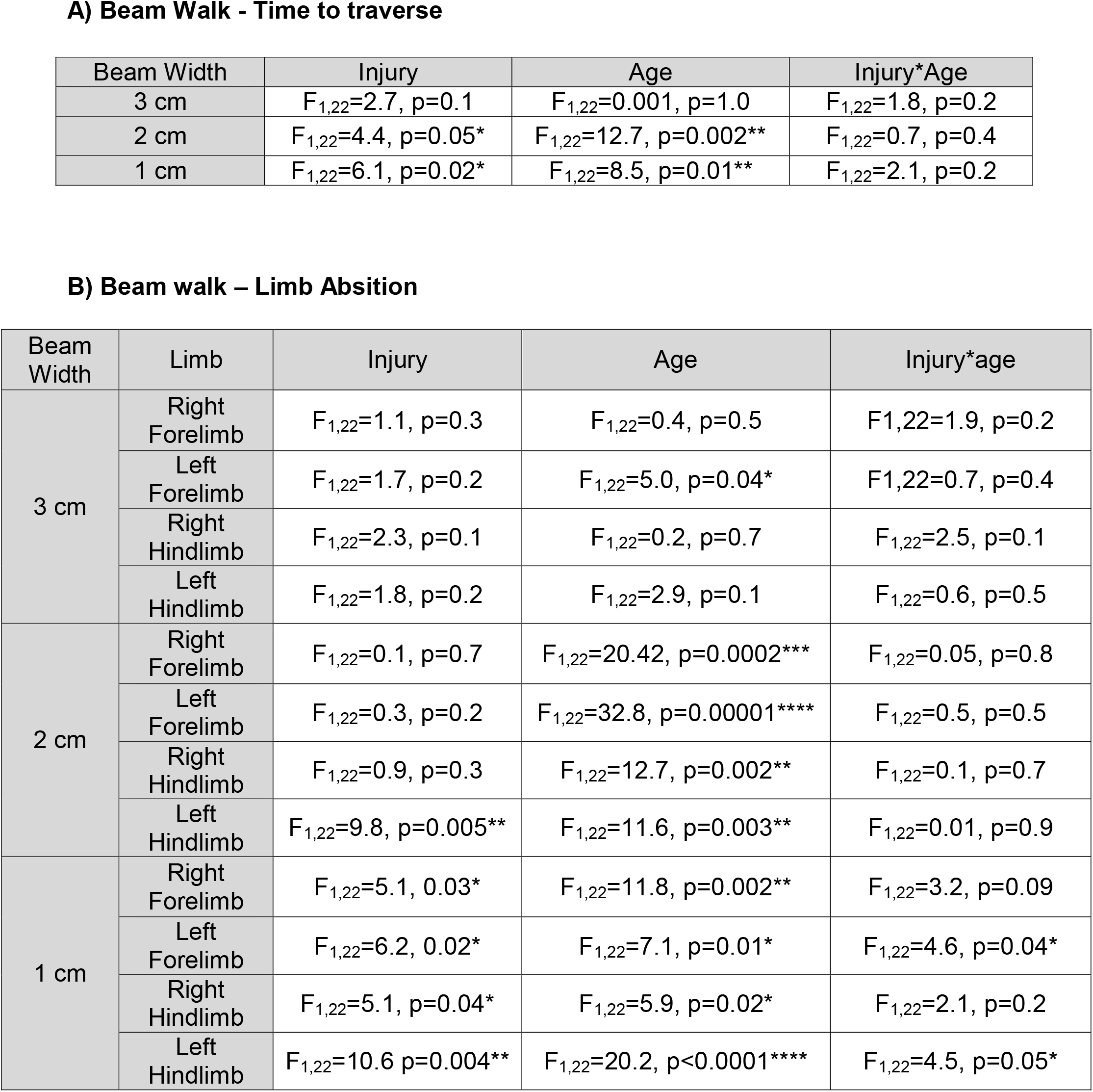
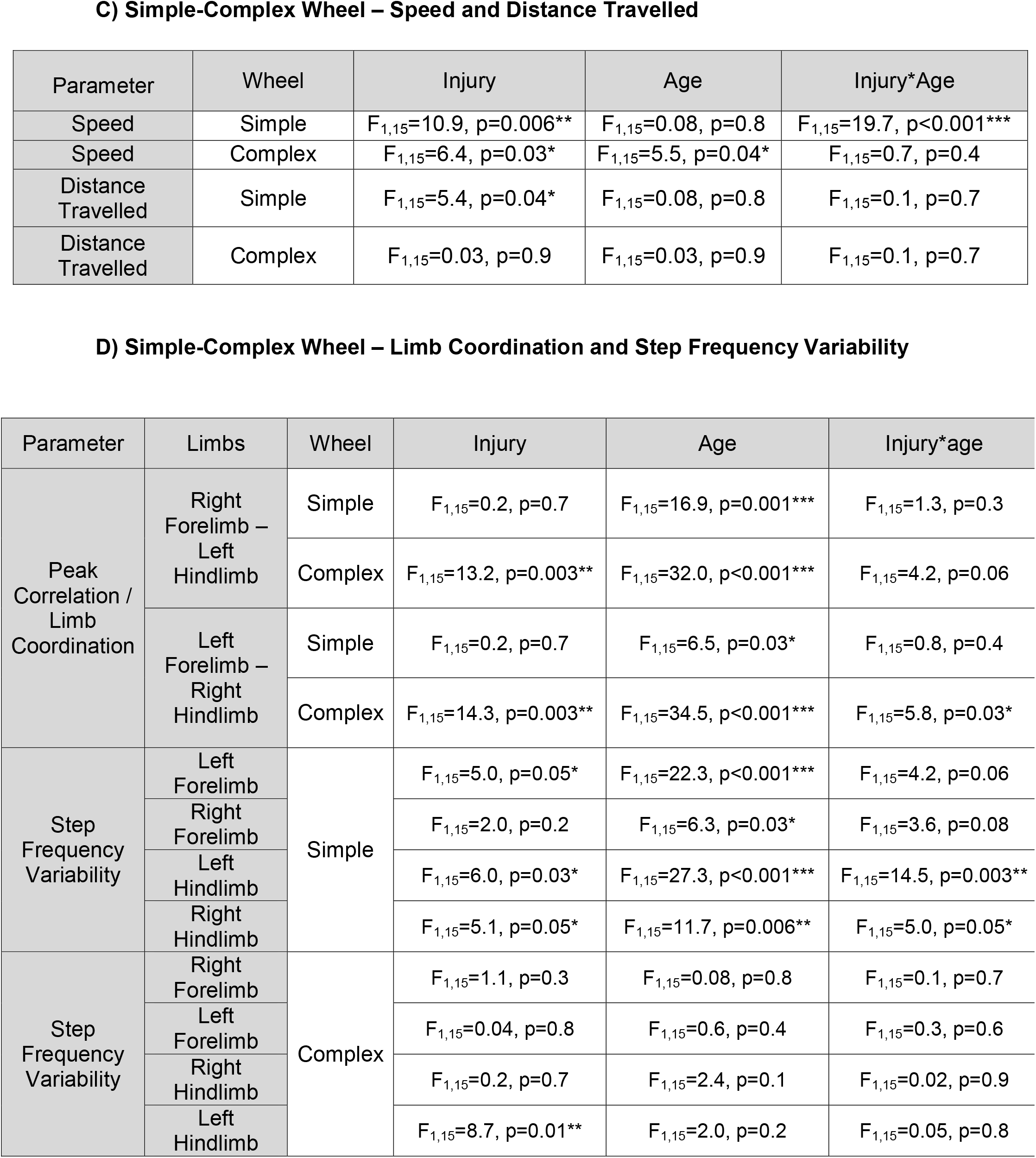
Two-way ANOVAs for Beam Walk and Simple Complex Wheel.

**A) Beam Walk - Time to traverse**
| Beam Width | Injury | Age | Injury*Age |
| --- | --- | --- | --- |
| 3 cm | $F_{1,22}=2.7, p=0.1$ | $F_{1,22}=0.001, p=1.0$ | $F_{1,22}=1.8, p=0.2$ |
| 2 cm | $F_{1,22}=4.4, p=0.05^*$ | $F_{1,22}=12.7, p=0.002^{**}$ | $F_{1,22}=0.7, p=0.4$ |
| 1 cm | $F_{1,22}=6.1, p=0.02^*$ | $F_{1,22}=8.5, p=0.01^{**}$ | $F_{1,22}=2.1, p=0.2$ |

**B) Beam walk – Limb Absition**
| Beam Width | Limb | Injury | Age | Injury*age |
| --- | --- | --- | --- | --- |
| 3 cm | Right Forelimb | $F_{1,22}=1.1, p=0.3$ | $F_{1,22}=0.4, p=0.5$ | $F_{1,22}=1.9, p=0.2$ |
| | Left Forelimb | $F_{1,22}=1.7, p=0.2$ | $F_{1,22}=5.0, p=0.04^*$ | $F_{1,22}=0.7, p=0.4$ |
| | Right Hindlimb | $F_{1,22}=2.3, p=0.1$ | $F_{1,22}=0.2, p=0.7$ | $F_{1,22}=2.5, p=0.1$ |
| | Left Hindlimb | $F_{1,22}=1.8, p=0.2$ | $F_{1,22}=2.9, p=0.1$ | $F_{1,22}=0.6, p=0.5$ |
| 2 cm | Right Forelimb | $F_{1,22}=0.1, p=0.7$ | $F_{1,22}=20.42, p=0.0002^{***}$ | $F_{1,22}=0.05, p=0.8$ |
| | Left Forelimb | $F_{1,22}=0.3, p=0.2$ | $F_{1,22}=32.8, p=0.00001^{****}$ | $F_{1,22}=0.5, p=0.5$ |
| | Right Hindlimb | $F_{1,22}=0.9, p=0.3$ | $F_{1,22}=12.7, p=0.002^{**}$ | $F_{1,22}=0.1, p=0.7$ |
| | Left Hindlimb | $F_{1,22}=9.8, p=0.005^{**}$ | $F_{1,22}=11.6, p=0.003^{**}$ | $F_{1,22}=0.01, p=0.9$ |
| 1 cm | Right Forelimb | $F_{1,22}=5.1, 0.03^*$ | $F_{1,22}=11.8, p=0.002^{**}$ | $F_{1,22}=3.2, p=0.09$ |
| | Left Forelimb | $F_{1,22}=6.2, 0.02^*$ | $F_{1,22}=7.1, p=0.01^*$ | $F_{1,22}=4.6, p=0.04^*$ |
| | Right Hindlimb | $F_{1,22}=5.1, p=0.04^*$ | $F_{1,22}=5.9, p=0.02^*$ | $F_{1,22}=2.1, p=0.2$ |
| | Left Hindlimb | $F_{1,22}=10.6 p=0.004^{**}$ | $F_{1,22}=20.2, p<0.0001^{****}$ | $F_{1,22}=4.5, p=0.05^*$ |

C) Simple-Complex Wheel – Speed and Distance Travelled
| Parameter | Wheel | Injury | Age | Injury*Age |
| --- | --- | --- | --- | --- |
| Speed | Simple | $F_{1,15}=10.9, p=0.006^{**}$ | $F_{1,15}=0.08, p=0.8$ | $F_{1,15}=19.7, p<0.001^{***}$ |
| Speed | Complex | $F_{1,15}=6.4, p=0.03^*$ | $F_{1,15}=5.5, p=0.04^*$ | $F_{1,15}=0.7, p=0.4$ |
| Distance Travelled | Simple | $F_{1,15}=5.4, p=0.04^*$ | $F_{1,15}=0.08, p=0.8$ | $F_{1,15}=0.1, p=0.7$ |
| Distance Travelled | Complex | $F_{1,15}=0.03, p=0.9$ | $F_{1,15}=0.03, p=0.9$ | $F_{1,15}=0.1, p=0.7$ |

D) Simple-Complex Wheel – Limb Coordination and Step Frequency Variability
| Parameter | Limbs | Wheel | Injury | Age | Injury*age |
| --- | --- | --- | --- | --- | --- |
| Peak Correlation / Limb Coordination | Right Forelimb – Left Hindlimb | Simple | $F_{1,15}=0.2, p=0.7$ | $F_{1,15}=16.9, p=0.001^{***}$ | $F_{1,15}=1.3, p=0.3$ |
| | | Complex | $F_{1,15}=13.2, p=0.003^{**}$ | $F_{1,15}=32.0, p<0.001^{***}$ | $F_{1,15}=4.2, p=0.06$ |
| | Left Forelimb – Right Hindlimb | Simple | $F_{1,15}=0.2, p=0.7$ | $F_{1,15}=6.5, p=0.03^*$ | $F_{1,15}=0.8, p=0.4$ |
| | | Complex | $F_{1,15}=14.3, p=0.003^{**}$ | $F_{1,15}=34.5, p<0.001^{***}$ | $F_{1,15}=5.8, p=0.03^*$ |
| Step Frequency Variability | Left Forelimb | Simple | $F_{1,15}=5.0, p=0.05^*$ | $F_{1,15}=22.3, p<0.001^{***}$ | $F_{1,15}=4.2, p=0.06$ |
| | Right Forelimb | | $F_{1,15}=2.0, p=0.2$ | $F_{1,15}=6.3, p=0.03^*$ | $F_{1,15}=3.6, p=0.08$ |
| | Left Hindlimb | | $F_{1,15}=6.0, p=0.03^*$ | $F_{1,15}=27.3, p<0.001^{***}$ | $F_{1,15}=14.5, p=0.003^{**}$ |
| | Right Hindlimb | | $F_{1,15}=5.1, p=0.05^*$ | $F_{1,15}=11.7, p=0.006^{**}$ | $F_{1,15}=5.0, p=0.05^*$ |
| Step Frequency Variability | Right Forelimb | Complex | $F_{1,15}=1.1, p=0.3$ | $F_{1,15}=0.08, p=0.8$ | $F_{1,15}=0.1, p=0.7$ |
| | Left Forelimb | | $F_{1,15}=0.04, p=0.8$ | $F_{1,15}=0.6, p=0.4$ | $F_{1,15}=0.3, p=0.6$ |
| | Right Hindlimb | | $F_{1,15}=0.2, p=0.7$ | $F_{1,15}=2.4, p=0.1$ | $F_{1,15}=0.02, p=0.9$ |
| | Left Hindlimb | | $F_{1,15}=8.7, p=0.01^{**}$ | $F_{1,15}=2.0, p=0.2$ | $F_{1,15}=0.05, p=0.8$ |

Forelimb absition on the 3cm beam shows a significant effect of age but no effect of injury or injury*age (Table 2B, Figure 2, Panel A). Biologically insignificant comparisons are responsible for age effects of left forelimb (F_3,22_=3.1, p=0.05). Hindlimb absition lacks significant effects of injury, age or injury*age (Table 2B,Figure 2, Panel A).

**Figure 2.**
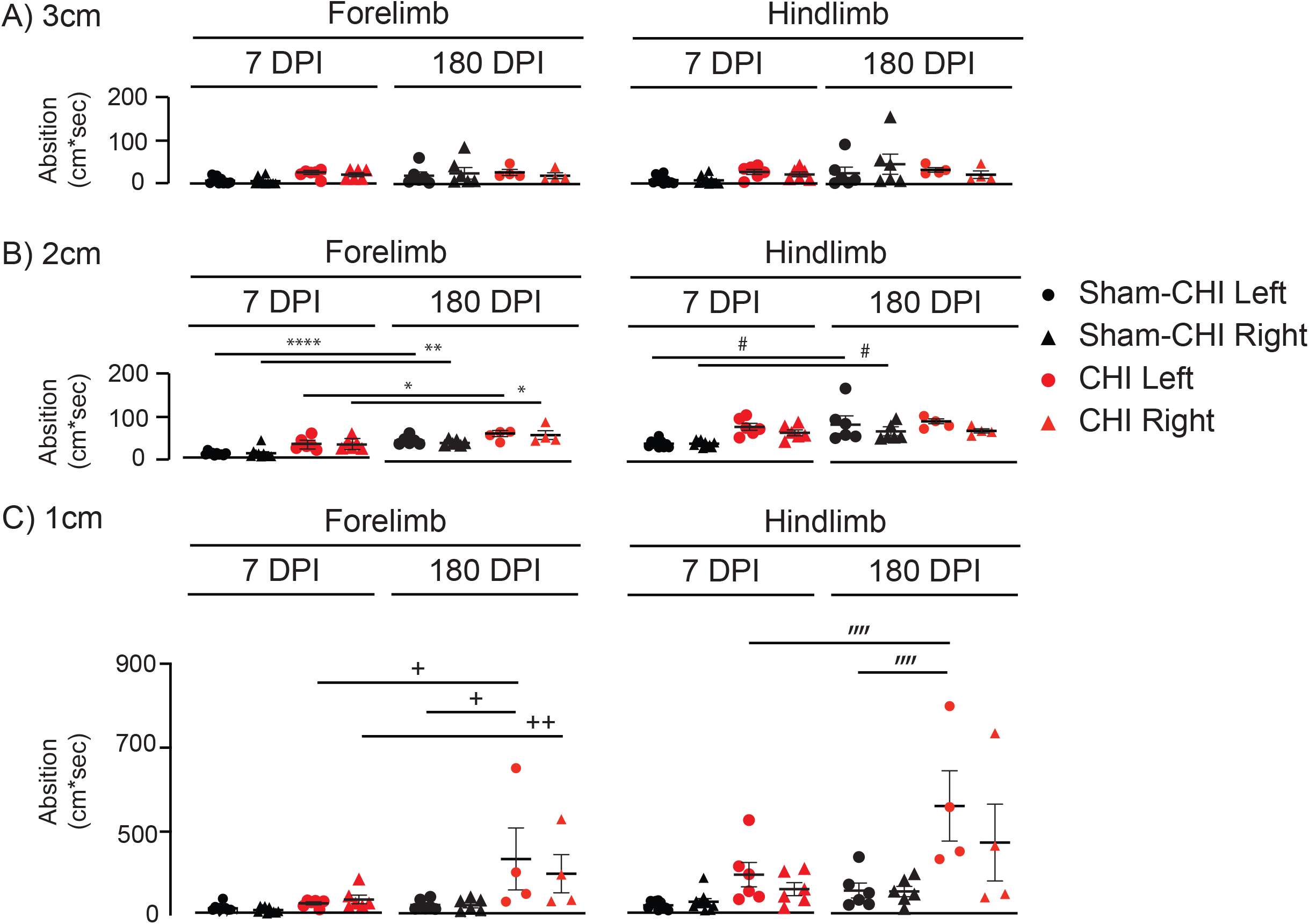
Beam Walk - Absition, **Panel A), 3cm,** No significant differences were seen on the 3cm beam. **Panel B), 2cm**, Both forelimbs of sham-CHI mice significantly differed between 7 and 180DPI for (Left, p<0.0001****; Right, p=0.005**) and CHI mice (Left, p=0.02* Right, p=0.05*). Both hindlimbs of sham-CHI mice significantly differed between 7 and 180 DPI (Left, p=0.05^#^; Right, p=0.02^#^). **Panel C), 1cm**, Both forelimbs of CHI mice significantly differed between 7 and 180 DPI. (Left, p=0.03^+^, Right, p=0.01^++^). The left forelimb at 180DPI significantly differed between sham-CHI and CHI mice.(p=0.03^+^). The left hindlimb of CHI mice significantly differed between 7 and 180 DPI (p=0.002^”“^). The left hindlimb at 180DPI significantly differed between sham-CHI and CHI mice (p=0.01^”“^).

On the 2cm beam absition of both forelimbs have a significant age effect, but no effects of injury or injury*age (Table 2B; Figure 2, Panel B). The left and right forelimbs of Sham-CHI and CHI mice have significantly increased absition compared to 7DPI (Left, F_3,22_=12.6, p<0.0001; Right, F_3,22_=7.3, p=0.001). Absition of the left hindlimb has a significant injury effect; both hindlimbs have a significant effect of age, yet no effect of injury*age. Injury trends to increase hindlimb absition in CHI mice at 7DPI (Left, F_3,22_=7.8, p=0.001, Sham vs CHI, p=0.07). Hindlimb absition in sham-CHI mice has a significant age effect (Left, p=0.05; Right, F_3,22_=5.1, p=0.008; p=0.005). These data suggest CHI mice have increased absition of the left hindlimb at 7DPI and age of sham mice increases absition of both hindlimbs (Table 2B; Figure 2, Panel B).

Absition of both forelimbs on the 1cm beam has significant injury and age effects (Table 2B; Figure 2, Panel C) with the left forelimb having a significant injury*age effect. Absition of both forelimbs of CHI mice has a significant group effect (Left, F_3,22_=4.8, p=0.01; Right, F_3,22_=5.6, p=0.005). Left forelimb absition increases in CHI mice between 7 and 180DPI (p=0.03). At 180 DPI, left forelimb absition of CHI mice is significantly greater than sham-CHI mice (Left: p=0.03; Right: p=0.07). These data suggest that a chronic and progressive increase in forelimb absition is produced by CHI. Hindlimb absition shows significant effects of injury and age, with the left hindlimb having a significant effect of injury*age (Table 2B; Figure 2, Panel C). Absition of both hindlimbs has a significant group effect (Left, F_3,22_=10.1, p<0.0001; Right, F_3,22_=3.7, p=0.03). Left hindlimb absition CHI mice at 180DPI was significantly larger than CHI mice at 7DPI (p=0.002) or Sham-CHI mice at 180DPI (p=0.01) (Supplementary Video 1). Right hindlimb absition of CHI mice at 180 DPI was larger than at 7DPI (p=0.09). These data suggest that injury produces a chronic and progressive increase in absition deficits in both forelimbs and the left hindlimb on the 1cm beam (Supplementary Video 1).

Speed on day 3 of simple wheel has a significant effect of injury and injury*age, but not age (Table 2C; Figure 3, Panel A). Speed on simple wheel has a significant group effect (F_3,15_=10.2, p=0.001). Sham mice run faster than injured mice at 14, but not 180 DPI (14 DPI, p<0.001; 180 DPI, p=0.9). Injured mice ran faster at 180DPI than 14DPI (p=0.03). Speed on day 3 of complex wheel running shows significant effects of injury, time, but not injury*time (Table 2C; Figure 3, Panel A). Speed on complex wheel had a significant group effect (F_3,15_=4.2, p=0.03) yet there were no biological significant pairwise differences. On simple wheel, distance traveled has a significant effect of injury but not age nor injury*age (Table 2C). There was no significant effect of group (F_3,15_=1.9, p=0.2). Distance traveled on complex wheel, had no significant effects of injury, age or injury*age (Table 2C). These data suggest that injury nor age affect the distance travelled on simple or complex wheel running.

**Figure 3.**
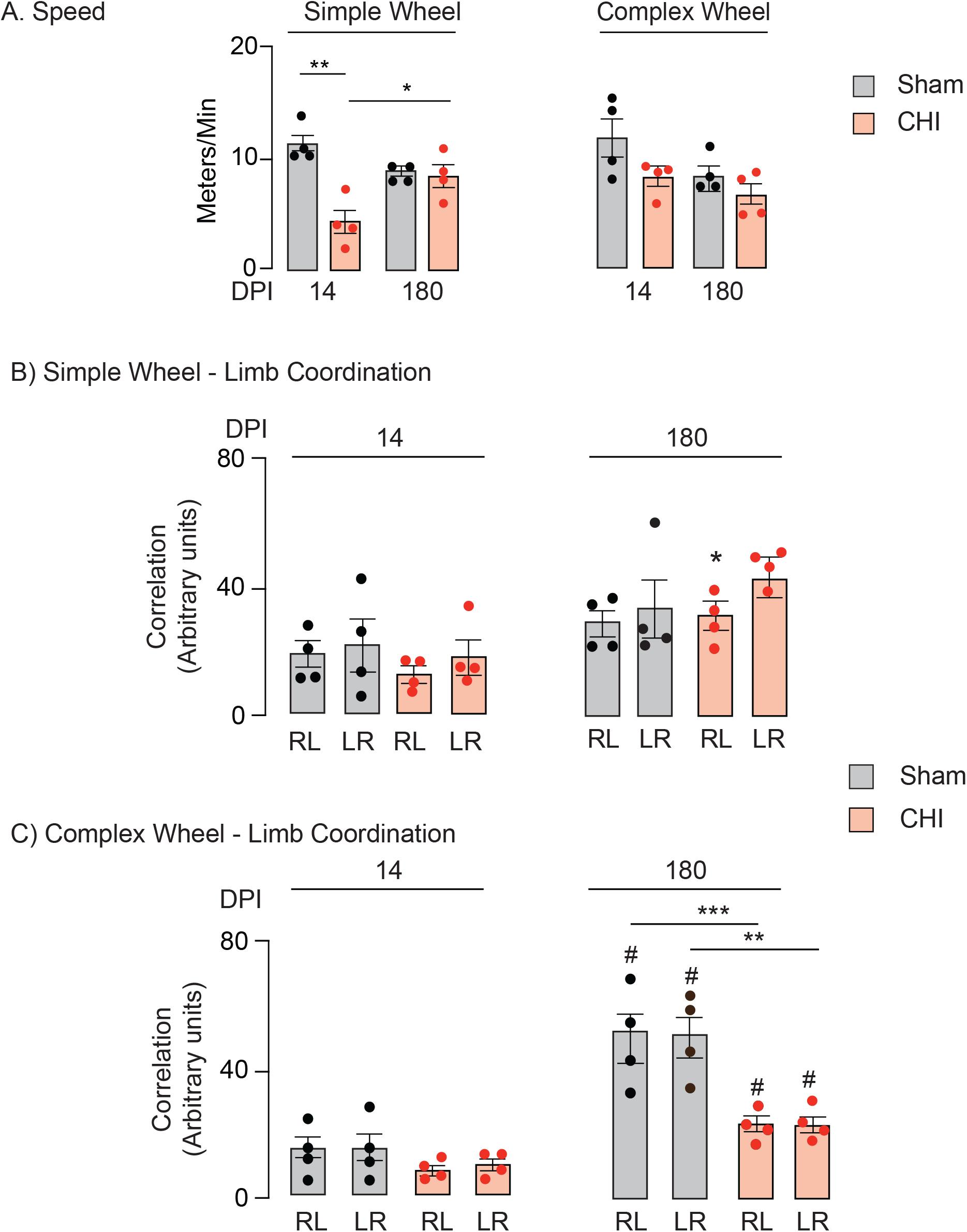
Simple-Complex Wheel – Speed and Limb Coordination. **Panel A) Speed** Average speed was assessed on simple-complex wheel. On simple wheel at 14DPI, sham-CHI mice ran faster than CHI mice (p<0.001***), CHI mice ran faster at 180DPI than 14DPI (p=0.03*). No biologically significant differences were observed on complex wheel. **Panels B and C) Limb Coordination** Right forelimb-left hindlimb (RL) and left forelimb-right hindlimb (LR) are assessed separately at 14 or 180 DPI. **Panel B), Simple Wheel**, RL coordination in sham-CHI and CHI mice at 180 DPI was significantly higher than CHI mice at 14 DPI (p=0.03*; p=0.01**). **Panel C), Complex Wheel**, RL and LR coordination of both sham-CHI and CHI mice was significantly higher at 180 DPI than at 14 DPI (RL: Sham-CHI14-Sham-CHI180, p<0.001***; Sham-CHI180-CHI14, P<0.001***; LR: Sham-CHI14-Sham-CHI180, p<0.001***; Sham-CHI180-CHI14, p<0.001***). RL and LR coordination was significantly lower in CHI mice than sham-CHI mice at 180DPI (RL: p=0.008**; LR: p=0.004**)

Speed but not distance travelled revealed deficits early after injury. However, neither outcome had persistent, delayed, or progressive deficits despite visible impairment. Therefore, DeepLabCut™ was employed to assess limb coordination on simple and complex wheel.

Limb coordination was assessed in the right forearm to the left hindlimb (RL) or the left forearm to the right hindlimb (LR) (Supplementary Video 2). RL and LR limb coordination on simple wheel had a significant effect of age but not injury or injury*age. RL but not LR limb coordination had significant group effects (RL, F_3,15_=6.1, p=0.009; LR, F_3,15_=2.5, p=0.1) (Table 2D; Figure 3, Panel B). RL coordination in injured mice increased at 180DPI compared to 14DPI (p=0.01). These data suggest that, on simple wheel, injury produces a mild change in RL limb coordination running that resolves over time. On complex wheel, both RL and LR coordination had significant effects of injury, age and injury*age. Both RL and LR had significant group effects (RL, F_3,15_=16.5, p<0.001; LR, F_3,15_=18.2, p<0.001) (Table 2D; Figure 3, Panel B). Between 14 and 180 DPI, coordination of both limb pairs increased in both sham (LR, p<0.001; RL, p<0.001) and injured mice (LR, p=0.008; RL, p=0.004). At 180, but not 14 DPI, both RL and LR coordination worsened between sham and injured mice (RL, p=0.008; LR, p=0.004). These data suggest a delayed and progressive decrease in the coordination of both limb pairs in injured mice on complex wheel (Supplementary Video 2). Injured mice had bilateral limb coordination deficits at 180 DPI yet showed no deficits in the more commonly assessed motor factors of speed or distance traveled. These data suggest that an altered form of limb coordination in injured mice compensates to maintain speed.

Impaired limb coordination in injured mice was further analyzed by examining each limb for the average step frequency and variability on simple and complex wheel. On simple wheel, both left and right forelimb step frequency showed an effect of injury and age, but not injury*age. Left forelimb showed a group effect (F_3,15_=3.4, p=0.05) yet pairwise comparisons showed no significant biological effects. Step frequency of the right hindlimb had a significant effect of injury and age but not injury*age and no significant group effects. Step frequency of the left hindlimb had significant effects of age but not injury or injury*age. The left hindlimb of injured mice showed a significant group effect (F_3,15_=7.1, p=0.005) with a lower frequency at 180 as compared to 14 DPI (p=0.007). Step frequency of the right hindlimb had significant effects of injury, but not age nor injury*age. The right hindlimb shows a significant group effect (F_3,15_=3.6, p=0.05) but no significant biological effects. Step frequency variability of the left forelimb showed significant effects of injury with both forelimbs showing significant effects of age but not injury*age (Table 2 D, Figure 4, Panel A). Both forelimbs had significant group effects (left, F_3,15_ =11.4, p=0.001; right, F_3,15_ =4.3, p=0.03). Step frequency variability of both forelimbs of injured mice differed between 14 and 180 DPI (left, p=0.002; right p=0.03). Hindlimb step frequency variability of both limbs had significant effects of injury, age and injury*age with significant group effects (left, F_3,15_ =17.2, p<0.001; right, F_3,15_ =7.8, p=0.004) with significant increase in variability in the injured group at 180 DPI compared to sham (left, p=0.006; right, p=0.05) or injured at 14DPI (left, p<0.001; right, p=0.007) (Table 2 D, Figure 4, Panel A). These data suggest injury produces a delayed increase in step frequency variability on simple wheel running. For complex wheel, average step frequency and step frequency variability for both fore and hindlimbs had no significant effects for injury, age or injury*age apart from the right hindlimb which has a significant injury effect (Table 2 D, Figure 4, Panel B). There were no biologically significant group effects. These data suggest that the step frequency of injured animals is altered on simple wheel, but not on complex wheel.

**Figure 4.**
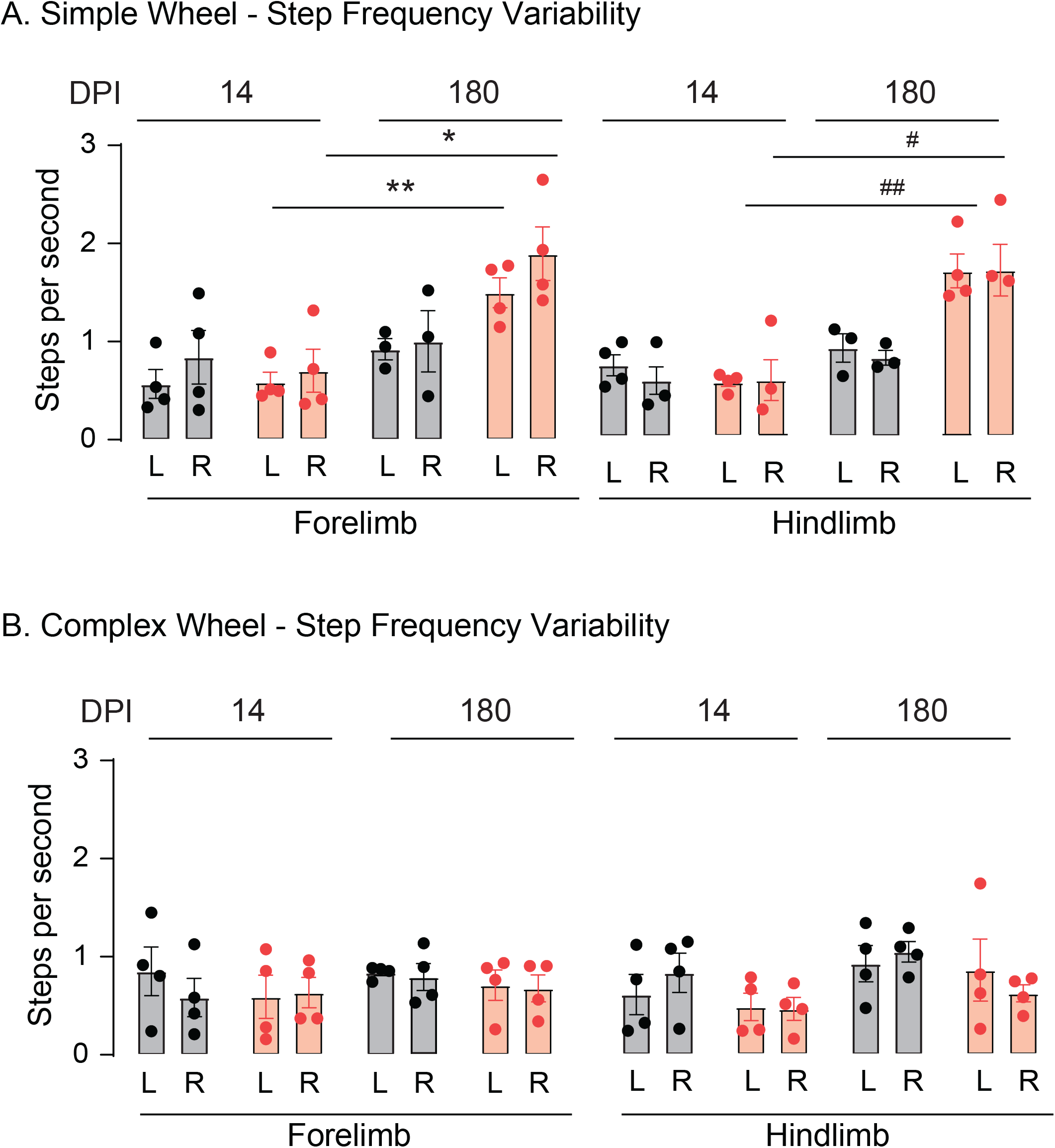
Simple-Complex Wheel – Step Frequency Variability, **Panel A), Simple Wheel** At 180 DPI, step frequency variability of the left and right forelimbs and hindlimbs of injured mice is significantly higher at 180 DPI than 14 DPI (Left forelimb, p=0.002**; Right forelimb, p=0.03*; Left hindlimb, p<0.001***; Right hindlimb, p=0.007**). Step frequency variability of both hindlimbs of injured mice is higher than sham mice at 180DPI (Left hindlimb, p=0.006**; Right hindlimb, p=0.05*). **Panel B, Complex Wheel** Step frequency variability did not differ significantly among groups.

Discussion

Chronic and progressive cognitive and motor deficits develop in one-third of patients who suffer a moderate to severe TBI(Foundas, 2013; Walker & Pickett, 2007; Wilson et al., 2017). Over time, patients with motor deficits may adopt compensatory strategies to perform complex tasks(Martini et al., 2011; Vasudevan et al., 2014). In animal models of TBI, however, chronic and progressive motor deficits are less commonly seen given the difficulty in distinguishing between effects of age and evolving injury, and reportedly normal function from compensatory strategies(Shultz et al., 2020; Xiong et al., 2013). The advent of customizable motor tracking in animals has thus provided a means to better distinguish these effects.

The CHI mouse model of TBI utilized in this study has shown both age and injury-dependent motor deficits both early and late after injury. Latency on rotarod shows a significant age effect and acute injury effect only at 7DPI (Figure 1, Panel C). Chronic and progressive deficits on open field, beam walk, and simple-complex wheel outcomes showed significant injury*age effects (Figures 1-4). Thus, a study of chronic and progressive deficits in preclinical TBI models must assess how much of a lack of deficit is due to the significant effects of aging in mice.

Injured mice at 180DPI significantly increased time spent in the open field arena center suggesting either lowered basal anxiety or somatomotor dysfunction (Figure 1, Panel A). Mice have a similar reduction in basal anxiety 7 weeks after a single controlled cortical impact(Popovitz et al., 2019). In the center of the arena, injured mice showed a strong turning bias toward the injured cortex (Figure 1, Panel B). Turning bias is thought to be a consequence of unilateral brain damage(Park et al., 2014). A similar turning developed bias is reported in rats 2 to 6 weeks after a single controlled cortical impact(Ajao et al., 2012). Additional examples of turning bias have been reported in models of cerebral ischemia and Parkinsonian deficits(Hartman et al., 2009; Park et al., 2014; Venna et al., 2014; Wang et al., 2006).

At 180 DPI. rotarod latency is dominated by age effects that overcome the acute effects of injury seen at 7 DPI (Figure 1, Panel C). Mice are heavier at 180DPI than at 7DPI and heavier mice perform worse on rotarod. These data suggest that the increased weight of mice at 180DPI underlies the large age effect on rotarod latency(Van Meer & Raber, 2005). Thus, a large age effect prevents rotarod from being used as an effective measure to examine chronic injury effects.

Age effects seen on time to traverse the 2cm and 1cm, but not the 3cm beams suggest that tasks with higher motor demand may readily detect chronic injury effects (Table 2A). The assay of forelimb and hindlimb absition using DeepLabCut™ tracking reveals chronic and progressive motor deficits not seen by assessing time to traverse the beam or number of foot-faults (Figure 2, Supplementary Video 1). Foot fault assessment did not reveal significant effects (Table 1), yet absition analysis of the same mice revealed both age and injury effects (Figure 2). An increase in absition without changing foot fault number suggests the use of alternative compensatory motor strategies to recover from foot faults.

Speed and distance traveled are commonly used motor outcomes for assessing motor function in mice. At 14 DPI injured mice were slower on simple wheel compared to sham, despite no change in distance travelled. Given the proximity of simple wheel assessment early after injury (Simple Wheel Day 3 = ∼4DPI), injured mice continue to demonstrate general signs of malaise. These data suggest injury acutely affects speed on day 3 of simple wheel running (∼4DPI) that is recuperated to sham levels at 180DPI. This recuperation in running speed at 180DPI, however, is accompanied by a dramatic increase in step frequency variability without much effect on limb coordination. On complex wheel, however, while sham and injured mice produce similar speed and distance travelled on complex wheel running, injured aged mice did not transfer the compensatory step frequency variability utilized on simple wheel, resulting in impaired limb coordination. This alteration of wheel dependent running pattern, simple wheel step frequency variability vs complex wheel limb coordination, is highly suggestive of a delayed motor compensation in injured mice.

Radiological and histological assessment of animals utilized in this study demonstrate a single CHI acutely produces atrophy to cortical hindlimb/torso motor and sensory, retrosplenial, and association areas, corpus callosum, and thalamic nuclei in the ipsilesional hemisphere(Havlicek et al., 2023). At 180DPI, hyperphosphorylated tau containing oligodendrocytes increase in density ipsilesionally in corpus callosum and thalamic nuclei, followed by continued radiological atrophy in the contralesional corpus callosum(Havlicek et al., 2023). Decreased ability to recover from a foot-fault may increase absition without changing the number of foot-faults. Increasing absition during recovery from foot-faults may suggest impairments arising from the injured sensory and motor cortices or thalamic nuclei during tasks of high motor demand(Heindorf et al., 2018; Schönfeld et al., 2017).

Increased step frequency variability on simple wheel running may result as a compensatory strategy to maintain sham level speed and limb coordination. On complex wheel, however, this compensatory strategy disappears followed by a significant reduction in limb coordination. The development of simple wheel compensatory running strategy and drop in complex wheel limb coordination may suggest ongoing impairment resulting from progressive corpus callosum atrophy(Havlicek et al., 2023).

One caveat of this study is that separate cohorts of mice are assessed at each time point. Mice assayed at 180 DPI were either sham-injured or injured immediately prior to the closing of the authors’ laboratory due to the COVID-19 pandemic which prevented behavioral assessment soon after injury. Ongoing studies are assaying the same cohort at acute and chronic timepoints.

## Acknowledgments

We thank the assistance of Jessy Lauer, PhD, with DeepLabCut™.

## Author Contributions

S.L., E.N. and P.J.B. designed the experiments in this study. S.L., and E.N. performed these studies. S.L., C.K. and P.J.B. analyzed the results. S.L., E.N., C.K., D.H. and P.J.B. wrote the manuscript.

## Author Disclosure

S.L., C.K., D.H., E.N., and P.J.B. have no conflicts of interest to declare.

This study was supported by NS108190 to P.J.B.

## Notes

### Competing Interest Statement

The authors have declared no competing interest.

### Summary of Updates

Title revised. Authorship updated. Text revised to remove assessment of cortical area and include the analysis of a new task, simple-complex wheel. Methods, Results, and Conclusions updated to include simple-complex wheel and reference radiological and histological data recently published using the same animals involved in this study. Figure 1 updated, Figure 2 updated, Figure 3 and 4 added, Supplemental videos 1 and 2 added, Table 2 added.

## References

Ajao, D. O., Pop, V., Kamper, J. E., Adami, A., Rudobeck, E., Huang, L., Vlkolinsky, R., Hartman, R. E., Ashwal, S., & Obenaus, A. (2012). Traumatic brain injury in young rats leads to progressive behavioral deficits coincident with altered tissue properties in adulthood. Journal of neurotrauma, 29(11), 2060–2074.

Bishop, K. M., Kruyer, A., & Wahlsten, D. (1996). Agenesis of the corpus callosum and voluntary wheel running in mice. Psychobiology, 24(3), 187–194.

Carter, R. J., Morton, J., & Dunnett, S. B. (2001). Motor coordination and balance in rodents. Current protocols in neuroscience, 15(1), 8.12. 11-18.12. 14.

Dean, P. J., & Sterr, A. (2013). Long-term effects of mild traumatic brain injury on cognitive performance. Frontiers in human neuroscience, 7, 30.

Flierl, M. A., Stahel, P. F., Beauchamp, K. M., Morgan, S. J., Smith, W. R., & Shohami, E. (2009). Mouse closed head injury model induced by a weight-drop device. Nature protocols, 4(9), 1328–1337.

Foundas, A. L. (2013). Apraxia: neural mechanisms and functional recovery. Handbook of clinical neurology, 110, 335–345.

Fox, G. B., Fan, L., Levasseur, R. A., & Faden, A. I. (1998). Sustained sensory/motor and cognitive deficits with neuronal apoptosis following controlled cortical impact brain injury in the mouse. Journal of neurotrauma, 15(8), 599–614.

Grin’kina, N. M., Li, Y., Haber, M., Sangobowale, M., Nikulina, E., Le’Pre, C., El Sehamy, A. M., Dugue, R., Ho, J. S., & Bergold, P. J. (2016). Righting reflex predicts long-term histological and behavioral outcomes in a closed head model of traumatic brain injury. PLoS One, 11(9), e0161053.

Hartman, R., Lekic, T., Rojas, H., Tang, J., & Zhang, J. H. (2009). Assessing functional outcomes following intracerebral hemorrhage in rats. Brain research, 1280, 148–157.

Havlicek, D. F., Furhang, R., Nikulina, E., Smith-Salzberg, B., Lawless, S., Severin, S. A., Mallaboeva, S., Nayab, F., Seifert, A. C., & Crary, J. F. (2023). A single closed head injury in male adult mice induces chronic, progressive white matter atrophy and increased phospho-tau expressing oligodendrocytes. Experimental neurology, 359, 114241.

Heindorf, M., Arber, S., & Keller, G. B. (2018). Mouse motor cortex coordinates the behavioral response to unpredicted sensory feedback. Neuron, 99(5), 1040-1054. e1045.

Hibbits, N., Pannu, R., Wu, T. J., & Armstrong, R. C. (2009). Cuprizone demyelination of the corpus callosum in mice correlates with altered social interaction and impaired bilateral sensorimotor coordination. ASN neuro, 1(3), AN20090032.

Johansson, B., Berglund, P., & Rönnbäck, L. (2009). Mental fatigue and impaired information processing after mild and moderate traumatic brain injury. Brain injury, (13-14), 1027–1040.

Kumar, R., Husain, M., Gupta, R. K., Hasan, K. M., Haris, M., Agarwal, A. K., Pandey, C., & Narayana, P. A. (2009). Serial changes in the white matter diffusion tensor imaging metrics in moderate traumatic brain injury and correlation with neuro-cognitive function. Journal of neurotrauma, 26(4), 481–495.

Lundqvist, A., Alinder, J., & Rönnberg, J. (2008). Factors influencing driving 10 years after brain injury. Brain injury, 22(4), 295–304.

Luong, T. N., Carlisle, H. J., Southwell, A., & Patterson, P. H. (2011). Assessment of motor balance and coordination in mice using the balance beam. JoVE (Journal of Visualized Experiments)(49), e2376.

Martini, D. N., Sabin, M. J., DePesa, S. A., Leal, E. W., Negrete, T. N., Sosnoff, J. J., & Broglio, S. P. (2011). The chronic effects of concussion on gait. Archives of physical medicine and rehabilitation, 92(4), 585–589.

Mathis, A., Mamidanna, P., Cury, K. M., Abe, T., Murthy, V. N., Mathis, M. W., & Bethge, M. (2018). DeepLabCut: markerless pose estimation of user-defined body parts with deep learning. Nature neuroscience, 21(9), 1281–1289.

McKenzie, I. A., Ohayon, D., Li, H., Paes de Faria, J., Emery, B., Tohyama, K., & Richardson, W. D. (2014). Motor skill learning requires active central myelination. science, 346(6207), 318–322.

Neumann, M., Wang, Y., Kim, S., Hong, S. M., Jeng, L., Bilgen, M., & Liu, J. (2009). Assessing gait impairment following experimental traumatic brain injury in mice. Journal of neuroscience methods, 176(1), 34–44.

Niechwiej-Szwedo, E., Inness, E., Howe, J., Jaglal, S., McIlroy, W. E., & Verrier, M. (2007). Changes in gait variability during different challenges to mobility in patients with traumatic brain injury. Gait & Posture, 25(1), 70–77.

Niogi, S., Mukherjee, P., Ghajar, J., Johnson, C., Kolster, R., Sarkar, R., Lee, H., Meeker, M., Zimmerman, R., & Manley, G. (2008). Extent of microstructural white matter injury in postconcussive syndrome correlates with impaired cognitive reaction time: a 3T diffusion tensor imaging study of mild traumatic brain injury. American Journal of Neuroradiology, 29(5), 967–973.

Park, S.-Y., Marasini, S., Kim, G.-H., Ku, T., Choi, C., Park, M.-Y., Kim, E.-H., Lee, Y.-D., Suh-Kim, H., & Kim, S.-S. (2014). A method for generating a mouse model of stroke: evaluation of parameters for blood flow, behavior, and survival [corrected]. Experimental neurobiology, 23(1), 104–114.

Pischiutta, F., Micotti, E., Hay, J. R., Marongiu, I., Sammali, E., Tolomeo, D., Vegliante, G., Stocchetti, N., Forloni, G., & De Simoni, M.-G. (2018). Single severe traumatic brain injury produces progressive pathology with ongoing contralateral white matter damage one year after injury. Experimental neurology, 300, 167–178.

Popovitz, J., Mysore, S. P., & Adwanikar, H. (2019). Long-term effects of traumatic brain injury on anxiety-like behaviors in mice: behavioral and neural correlates. Frontiers in behavioral neuroscience, 13, 6.

Schönfeld, L.-M., Dooley, D., Jahanshahi, A., Temel, Y., & Hendrix, S. (2017). Evaluating rodent motor functions: Which tests to choose? Neuroscience & Biobehavioral Reviews, 83, 298–312.

Shultz, S. R., McDonald, S. J., Corrigan, F., Semple, B. D., Salberg, S., Zamani, A., Jones, N. C., & Mychasiuk, R. (2020). Clinical relevance of behavior testing in animal models of traumatic brain injury. Journal of neurotrauma, 37(22), 2381–2400.

Tsenter, J., Beni-Adani, L., Assaf, Y., Alexandrovich, A. G., Trembovler, V., & Shohami, E. (2008). Dynamic changes in the recovery after traumatic brain injury in mice: effect of injury severity on T2-weighted MRI abnormalities, and motor and cognitive functions. Journal of neurotrauma, 25(4), 324–333.

Van Meer, P., & Raber, J. (2005). Mouse behavioural analysis in systems biology. Biochemical Journal, 389(3), 593–610.

Vasudevan, E. V., Glass, R. N., & Packel, A. T. (2014). Effects of traumatic brain injury on locomotor adaptation. Journal of neurologic physical therapy, 38(3), 172–182.

Venna, V. R., Li, J., Hammond, M. D., Mancini, N. S., & McCullough, L. D. (2014). Chronic metformin treatment improves post-stroke angiogenesis and recovery after experimental stroke. European Journal of Neuroscience, 39(12), 2129–2138.

Walker, W. C., & Pickett, T. C. (2007). Motor impairment after severe traumatic brain injury: A longitudinal multicenter study. Journal of Rehabilitation Research & Development, 44(7).

Wang, Y., Galvan, V., Gorostiza, O., Ataie, M., Jin, K., & Greenberg, D. A. (2006). Vascular endothelial growth factor improves recovery of sensorimotor and cognitive deficits after focal cerebral ischemia in the rat. Brain research, 1115(1), 186–193.

Wilson, L., Stewart, W., Dams-O’Connor, K., Diaz-Arrastia, R., Horton, L., Menon, D. K., & Polinder, S. (2017). The chronic and evolving neurological consequences of traumatic brain injury. The Lancet Neurology, 16(10), 813–825.

Xiong, Y., Mahmood, A., & Chopp, M. (2013). Animal models of traumatic brain injury. Nature Reviews Neuroscience, 14(2), 128–142.

