## Supplementary Video 1 Legend for "Analysis of markerless limb tracking reveals chronic and progressive motor deficits after a single closed head injury in mice"

**Supplementary Video 1, Beam Walk –** DeepLabCut^TM^ **Tracking with Absition assessment.** Representative video recording and absition trace with of sham-CHI and CHI mice assessed at 180DPI on the 1cm beam. View is of the left side of the animal and has been flipped to correspond with absition trace direction. The top of the beam corresponds to 0 on the y-axis of the absition trace.
