## Supplementary Video 2 Legend for "Analysis of markerless limb tracking reveals chronic and progressive motor deficits after a single closed head injury in mice"

**Supplementary Video 2, Complex Wheel running Day 3 – DLC Tracking and Limb Coordination** Representative video recording with DeepLabCut^TM^ tracking and custom python script derived limb coordination trace of a sham-CHI and CHI mouse assessed at 180DPI on Day 3 of Complex Wheel running. View is of the left side of the animal and has been flipped to correspond with limb coordination trace direction. Y-axis is the distance from mouse midpoint to the end of each limb in pixels. X-axis is time running in seconds.
